# *De novo* genome Assembly of *Rauvolfia Serpentina* and comparative paleodemographic analysis of *Apocynaceae* plants

**DOI:** 10.1101/2025.10.27.684745

**Authors:** Mratunjay Dwivedi, Nagarjun Vijay

## Abstract

*Rauvolfia serpentina* is a perennial subshrub widely distributed across Asia and the Indian subcontinent. Belonging to the medicinally important *Apocynaceae* family, it is renowned for its ecological importance and ethnobotanical applications, especially through the biosynthesis of therapeutic indole alkaloids. In this study, we present a de novo genome assembly and annotation for *R. serpentina*. The assembled genome comprises approximately 184,000 scaffolds with a scaffold N50 of 7.18 Kbp and a high BUSCO completeness score of ∼90% encompassing ∼22,000 annotated genes. We analysed the paleodemographic histories of nine additional *Apocynaceae* species using Pairwise Sequentially Markovian Coalescent (PSMC) modelling to place these findings in a broader evolutionary context. This comparative analysis revealed a consistent signature of a pronounced bottleneck across the family during the Mid-Pleistocene glaciations, with few notable species showing high adaptive resilience marked by early recovery in effective population size (N_e_), while some even showed secondary N_e_ peaks in the warmer interglacial periods. Although marked by complex and fairly distinct demographic trajectories, most species exhibit stabilised N_e_ near the onset of the Holocene. The availability of this high-quality draft genome assembly provides an important resource to advance functional, ecological, and comparative genomics in *R. serpentina* as well as aid heterologous production of pharmaceutically important biomolecules at an industrial scale, thereby alleviating the pressure on wild populations.

## Introduction

*Rauvolfia serpentina* (commonly known as *Sarpgandha, Chandrika, Serpentine wood, Indian snakeroot*, or *Devil peppers*) is a perennial, evergreen subshrub belonging to the *Apocynaceae* family within the order *Gentianales*. Native to Southeast Asia, the species is now found widely distributed across tropical regions, including parts of Africa, America, Sri Lanka, Nepal, Myanmar, Malaysia, Indonesia, and most notably along the lower Gangetic plains and the sub-Himalayan belt of the Indian subcontinent (Mukherjee et al., 2019) where it has been historically recognised within the traditional systems of medicine, with ancient Ayurvedic texts such as the Sushruta Samhita, Charaka Samhita, Ashtanga Hridaya and Vrindamadhava dating back to around 1000 BCE documenting its medical usage (Lobay, 2015).

The pharmacological relevance of *R. serpentina* is primarily attributed to its diverse repertoire of secondary metabolites, particularly the monoterpene indole alkaloids (MIAs), which include reserpine, reserpiline, rescinnamine, ajmaline, ajmalacine, deserpidine, serpinine, serpentine, serpentinine, rauvolfinine, vomilenine, yohimbine, picrinine, norseredamine, vinorine, and seredamine, among others (Dey and De, 2010) (Pandey et al., 2016). In addition, other important phytochemicals such as stigmasterol (Dey et al., 2016; Dey and Pandey, 2014) and rutin (Dey and De, 2010) have also been reported.

These compounds have been reported to exhibit a broad spectrum of biological activities, including antimalarial and anticancer effects (Beljanski and Beljanski, 1986, 1984, 1982; Wright et al., 1996),(Bae et al., 2020; Lin et al., 2012) with demonstrated therapeutic potential in the treatment of cardiovascular disorders such as hypertension (Achor et al., 1955; Lobay, 2015) and arrhythmias (Brugada et al., 2003; Rolf et al., 2003; Wolpert et al., 2005), as well as in managing anxiety, epilepsy, trauma, and gastrointestinal disturbances (Meena et al., 2009; Poonam, 2013).

Alongside these well-documented applications, traditional ethnomedicinal uses of *R. serpentina* also encompass treatment for snakebite, insect stings, psychiatric conditions, including schizophrenia and insomnia, dermatological conditions, respiratory ailments, fever, pneumonia, body pain, scabies, spleen disorders, ocular diseases, and even AIDS and various veterinary diseases (Dey and De, 2011, 2010; Pandey et al., 2016). However, despite its well-established pharmacological significance and extensive biochemical profiling, *R. serpentina* remains relatively under-investigated at the genomic level.

Adequate genomic data could not only provide insight into its evolutionary lineage but also help reveal the molecular pathways underlying its biosynthetic capabilities. In this study, we perform high-coverage whole-genome sequencing of *R. serpentina* and report its first de novo genome assembly and annotation. Our study will not only expand the genomic repertoire of the medicinally valuable *Apocynaceae* family but also serve as a foundational resource for future molecular and biotechnological research, potentially expanding its therapeutic applications and aiding in the development of effective in vitro conservation and genetic improvement strategies for its sustainable utilisation and long-term preservation (Gantait et al., 2017).

## Methods

### Sample Collection and DNA Sequencing

Fresh leaves of *R. serpentina* were harvested from a shrub located within the premises of the Indian Institute of Science Education and Research, Bhopal (GPS coordinates: 23°17′18.8196″N, 77°16′29.946″E; altitude: 520 meters above sea level). Genomic DNA was isolated using the Qiagen DNeasy Kit. The quality and integrity of the extracted DNA were evaluated through 1× agarose gel electrophoresis, Nanodrop spectrophotometry, and Qubit fluorometry. Short-read sequencing libraries (150 bp) were prepared using the Kapa HyperPrep Kit with an average insert size of 250 ± 50 bp. Approximately 83 Gbp of paired-end sequencing data was generated using the Illumina NovaSeq 6000 platform.

The raw sequencing data quality was assessed using FASTQC version 2.3.0 (Andrews et al., 2015), and the results were aggregated and visualised using MULTIQC version 1.22.2 (Ewels et al., 2016). To estimate the genome size, a k-mer-based approach was employed using Jellyfish version 1.1.12 (Marçais and Kingsford, 2011), followed by modelling with GenomeScope version 1.0.0 (Vurture et al., 2017) using a k-mer length of 21.

### Genome Assembly

The genome of *R. serpentina* was assembled using Platanus version 1.2.4 (Kajitani et al., 2014), optimised for highly heterozygous plant genomes. Assembly parameters included an initial k-mer size of 35, a bubble crush threshold of 0.2 for resolving ambiguous paths, a minimum k-mer coverage of 10, and a k-mer extension step size of 10. The same pipeline was used for scaffold formation and gap filling to get the final genome assembly (see **Fig. 1**).

**Fig. 1.**
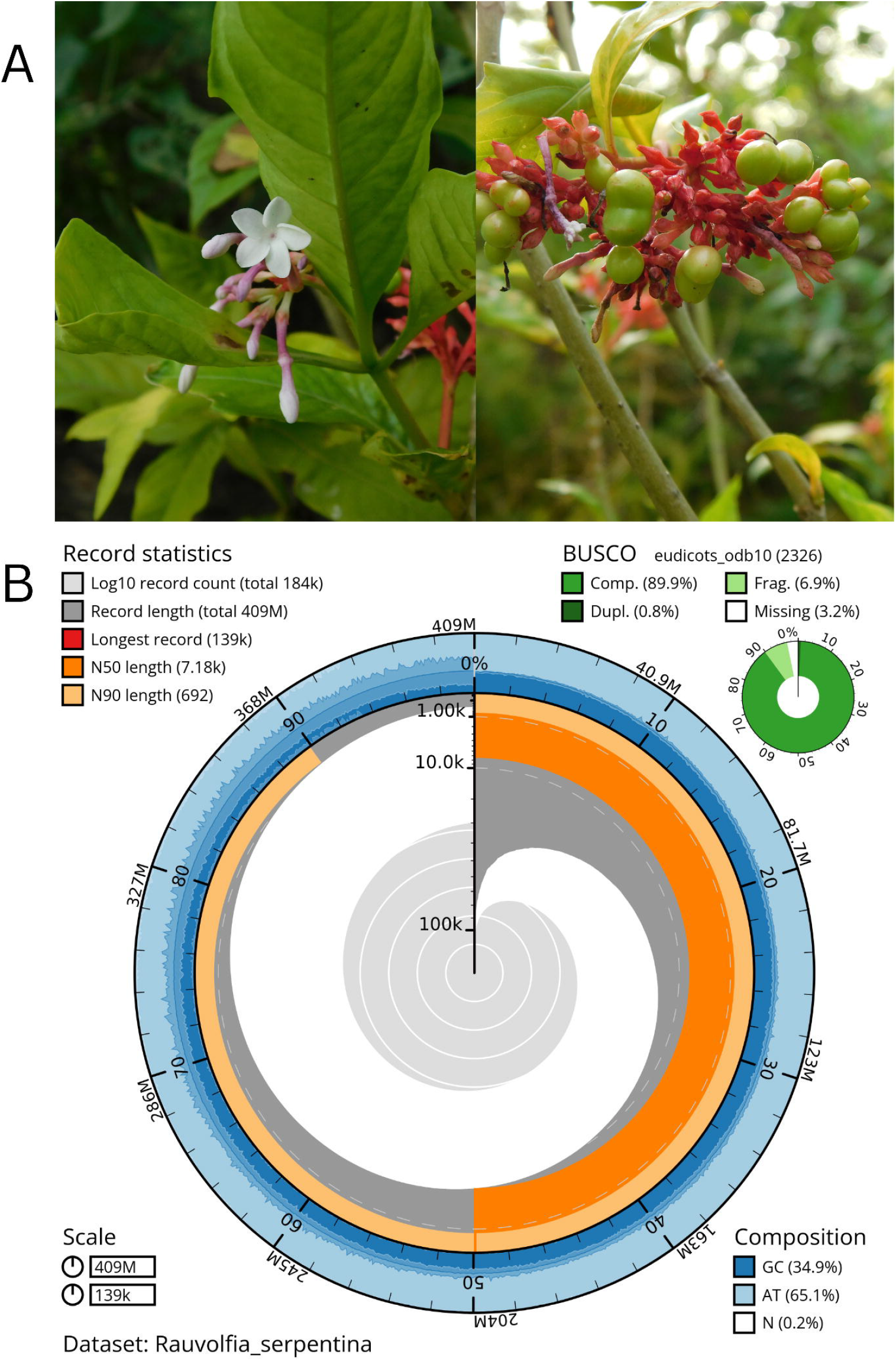
Snailplot summarising assembly metrics and gene completeness for the *Rauvolfia serpentina* genome. **(A)** *R. serpentina* is an important medicinal plant of the *Apocynaceae* family. **(B)** The dark grey area depicts the distribution of scaffold lengths, with the plot radius scaled to the longest scaffold assembled (139 kbp). Orange and pale orange segments indicate the N50 and N90 scaffold lengths, respectively. The pale grey spiral represents the cumulative scaffold count plotted on a logarithmic scale. The outer ring, shaded in blue and pale blue, illustrates the distribution of GC, AT, and ambiguous (N) base percentages. The green pie chart in the upper right corner shows the proportion of BUSCO genes identified as complete or missing.

Then, the quality of the assembled genome was comprehensively evaluated, and multiple metrics were employed to assess assembly continuity, completeness, and repeat content. QUAST version 5.2.2 (Mikheenko et al., 2018) was used to compute standard assembly statistics representing genome continuity, including N50, N75, longest contig length, and the number of ambiguous bases (N’s) per 100 Kbp (see **Supplementary Table S1**).

The completeness of the assembly was assessed using BUSCO version 5.8.2 (Simão et al., 2015), which quantifies the presence of conserved single-copy orthologous genes. Analyses were conducted using both the eudicotyledons_odb10 (see **Supplementary Table S2**) and embryophyta_odb10 (see **Supplementary Table S3**) lineage datasets, alongside comparisons to previously published genomes within the family *Apocynaceae*.

In addition to these structural metrics, the assembly quality in repetitive, low-complexity genomic regions was also evaluated through the LAI (LTR Assembly Index) score, calculated using the LAI module of LTR-retriever version 2.9. The repeats were further visualised using Repeat Landscape (Ou and Jiang, 2018) (see **Supplementary Table S4**).

### Genome Annotation

Structural gene annotation was performed using MAKER-P version 3.01.04 (Campbell et al., 2014) with MPI for parallel processing. Repeat libraries were generated using LTR-retriever version 2.9.0 (Ou and Jiang, 2018) and used to mask repetitive elements within the genome.

For the initial round of de novo gene prediction, the coding sequence (CDS) dataset of *Rauvolfia tetraphylla* (Lezin et al., 2024) and a concatenated multi-fasta of protein sequences of *Rauvolfia tetraphylla and Arabidopsis thaliana* were used as homology evidence. For further rounds of annotation, the results from previous annotation rounds progressively serve as the training set for gene prediction tools, SNAP version 2.46.1 (Korf, 2004) and AUGUSTUS version 3.3.3 (Stanke et al., 2008), implemented in BUSCO version 5.8.2 (Simão et al., 2015).

Annotation was iteratively refined across four rounds until convergence was achieved, as indicated by stabilisation of the Annotation Edit Distance (AED) values (see **Supplementary Table S5**).

### Organelle Genome Assembly and Annotation

The raw sequencing read data were used to assemble the chloroplast genome using the NOVOPlasty version 4.3.5 (Dierckxsens et al., 2017) program. The pbsA gene sequence of *R. serpentina* (Gene ID: 54600169) was used as a seed sequence. The assembler uses the seed sequence to find reads that cover this sequence and initiates overlapped sequence assembly. The assembled chloroplast genome consisted of one contig (see **Fig. 2A**). The full length of the assembled chloroplast genome was 155.3 Kbp. It was then annotated using GeSeq and visualised using OGDRAW implemented in CHLOROBOX (Greiner et al., 2019). For mitochondrial genome assembly, the *nad7* gene sequence of *Trachelospermum jasminoides* (Gene ID: 88599469) was used as a seed, and the *T. jasminoides* mitochondrial genome (Accession ID: OR333986.1) was used as a reference. The total assembled mitochondrial sequence was 627.3 Kbp (see **Fig. 2B**).

**Fig. 2.**
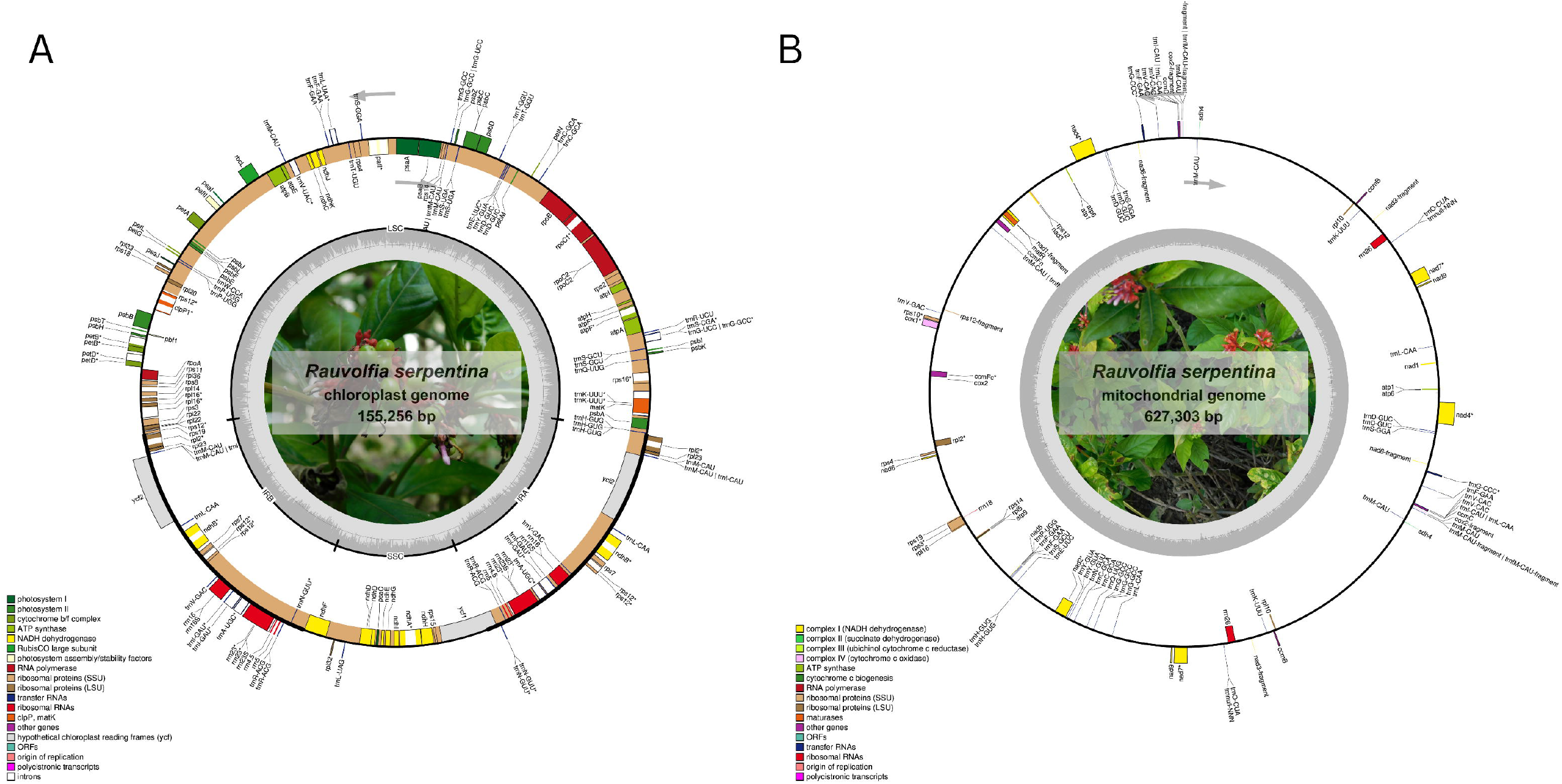
Chloroplast and mitochondrial genome maps of *Rauvolfia serpentina*. **(A)** Circular representation of the *R. serpentina* chloroplast genome. The inner circle highlights the major structural regions: the Large Single Copy (LSC), Small Single Copy (SSC), and the Inverted Repeats (IRA and IRB). The outermost ring shows gene positions and functional annotations, with gene classes colour-coded according to functional categories indicated in the legend (lower left). **(B)** Circular map of the *R. serpentina* mitochondrial genome. Genes transcribed on the positive strand are placed on the outer circle, while those on the negative strand appear on the inner circle. Genes containing introns are marked with an asterisk (*). Functional categories are colour-coded, and the transcription direction is clockwise (outer) or counterclockwise (inner). The innermost dark grey plot indicates the GC content across the mitochondrial genome.

### Phylogenetic analysis and genome comparison

For phylogenetic analysis, 22 complete chloroplast genomes from the *Apocynaceae* family were downloaded from NCBI, followed by alignment using MAFFT v7.525 (Katoh et al., 2002; Katoh and Standley, 2013). The phylogenetic tree was generated using maximum likelihood (ML) implemented in RaxML (Stamatakis, 2014). Phylogenetic robustness was evaluated using 1000 bootstrap replicates, and the resulting tree was visualised in Figtree (Rambaut, 2016, 2009). Additionally, the comparison of the borders of large single copy (LSC), small single copy (SSC), and inverted repeat regions (IR) in the chloroplast genome was performed with *Cynanchum wilfordii, Catharanthus roseus, R. serpentina, Asclepias syriaca and Rhazya stricta* using IR scope (Amiryousefi et al., 2018).

### Demographic history inference

Published plant genome assemblies of species belonging to the *Apocynaceae* family, particularly *Apocynum pictum, Apocynum venetum, Asclepias incarnata, A. syriaca, C. roseus, C. wilfordii, Echites panduratus, Marsdenia floribunda, and Pachypodium lamerei*, were downloaded from NCBI. The genomes and corresponding short-read data were downloaded from public repositories and mapped to the corresponding genomes using the minimap2 read mapper (Li, 2018). PSMC analysis (Li and Durbin, 2011) was performed on the majority of these species with default parameters, i.e., -t5 –r5 –p “4 + 25*2 + 4 + 6” while *C. wilfordii* and *A. venetum* were analysed with -p “10 + 18*2 + 6”. These parameter settings of PSMC were found to result in more than 10 recombination events in each atomic interval after 20 iterations, thereby enhancing the resolution and reliability of inferred historical effective population sizes. A uniform mutation rate estimate of 2.5e-09 per site per year previously established for *Populus trichocarpa* in an earlier study (Bai et al., 2018) was used for all the species. Species-specific per-generation mutation rates were obtained using corresponding generation times curated through a literature search (see **Supplementary Table S6**).

## Results

### Genome Assembly and Annotation

The *de novo* assembly of *R. serpentina* yielded a draft genome of length 408.6 Mb, distributed across 184,278 scaffolds, which was a little less than the genome size of 513.7 Mb estimated using GenomeScope (see **Supplementary Fig. 1**), through *k*-mer frequency analysis of the raw reads.

Assembly contiguity was assessed using standard metrics. The N50 scaffold length was 7.18 Kbp, and the L50 value was 11,573, while the longest scaffold assembled was 139 Kbp. Evaluation of repeat content using LTR-retriever yielded a LTR Assembly Index (LAI) score of 1.2, suggesting the assembly captured low-complexity repetitive regions, albeit with limited resolution in long terminal repeat (LTR) regions. The relative abundance of repeat classes was further visualised using Repeat Landscape (see **Supplementary Fig. 2**).

Assessment and comparison of genome completeness with other *Apocynaceae* genomes was performed using BUSCO v5.8.2, using both eudicotyledons_odb10 and embryophyta_odb10 datasets, reporting a comparable and complete genome with 89.9% and 90.6% complete gene sets present, respectively, in the datasets (see **Fig. 3**). Gene annotation predicted around 21,983 genes with an average length of 3636.31. These results collectively indicate a reasonably complete draft genome, suitable for downstream comparative analyses.

**Fig. 3.**
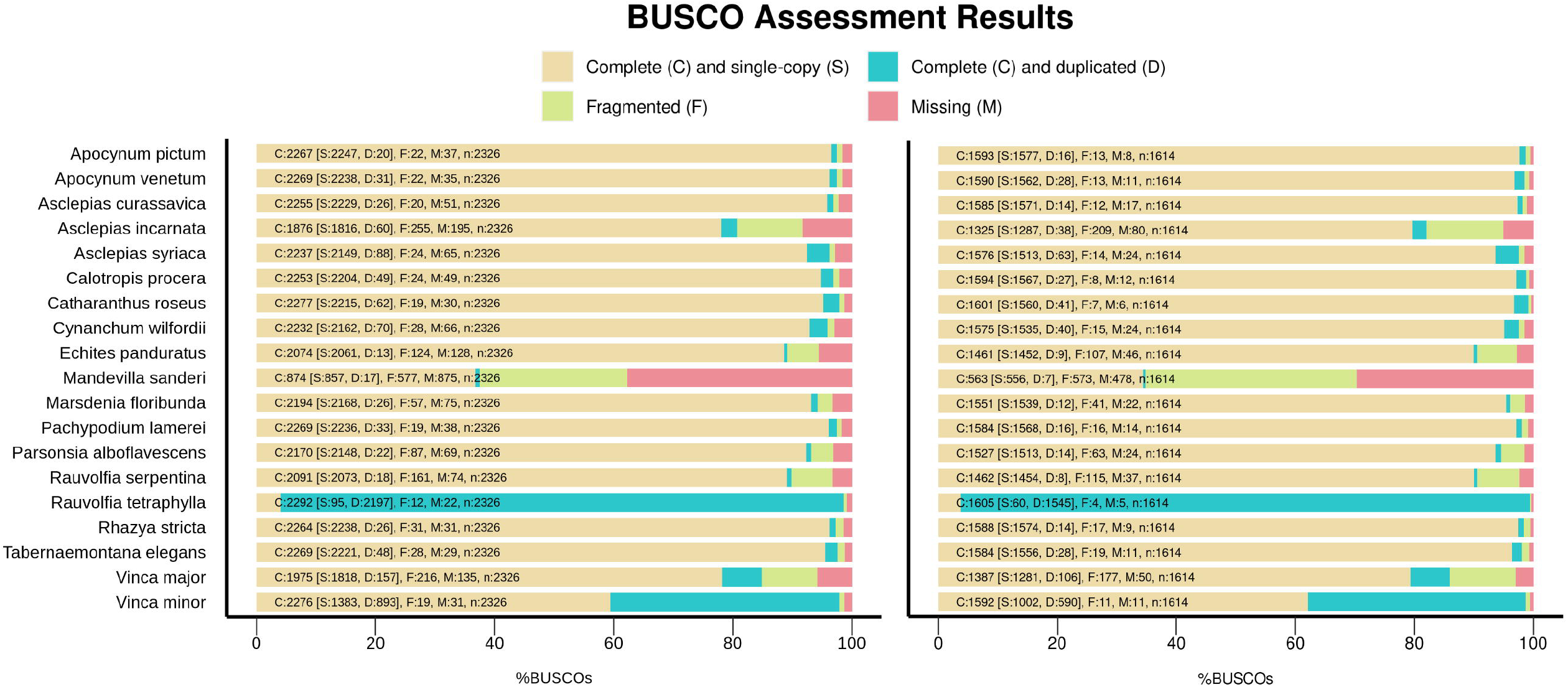
Assembly quality comparison of the *Apocynaceae* family using BUSCO. Assembly quality comparison of the *Apocynaceae* family using the Eudicotyledons_odb10 (left) and Embryophyta_odb10 (right) dataset. Comparison of genome completeness based on BUSCO scores of *Apocynaceae* genome assemblies. *R. serpentina* showed relatively complete assembly compared to other species.

### Phylogenetic analysis and genome comparison

The evolutionary placement of *R. serpentina* was determined using the chloroplast genomes of 22 other species of the *Apocynaceae* family to generate the phylogenetic tree. The presented chloroplast assembly was found to cluster with other species in the *Rauvolfia* family in the generated phylogenetic tree (see **Supplementary Fig. 3**). The LSC, SSC, and IR areas of *R. serpentina* relative to other *Apocynaceae* chloroplast genomes were visualised to evaluate differences in size and junction (see **Supplementary Fig. 4**).

### Demographic Analysis

PSMC plots estimated highly variable N_e_ curves from ∼10 million years ago to 10 KYA (based on scaling parameters) (see **Fig. 4**). All the species exhibit at least one cycle of demographic expansion followed by contraction. The curves show strong increases in N_e_ at ∼5 Ma, with N_e_ peaks in the mid-Pleistocene period, following an immediate decline and stabilisation at ∼50 KYA. Other than these limited occurrences of similarity, we observed a complex demographic history in *Apocynaceae* species with lineages following varying trends of expansion and contraction through time.

**Fig. 4.**
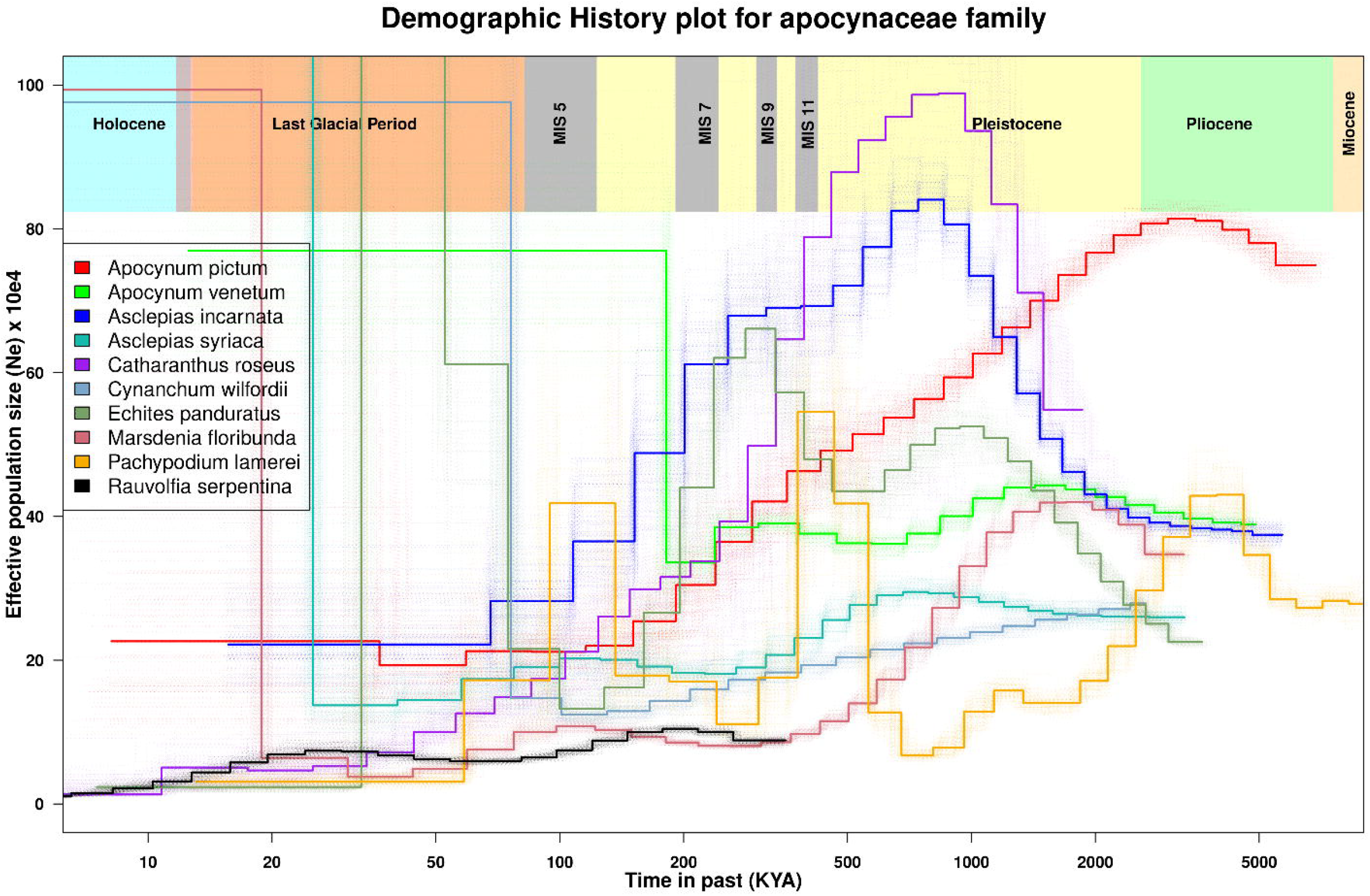
Comparative demographic histories of *Apocynaceae* plants, PSMC-inferred *Ne* trajectories for ten *Apocynaceae* species are shown with bootstrap support. Rectangles at the top indicate time intervals corresponding to major predicted glaciation events. *R. serpentina* displays a notably stabilised *Ne* trajectory compared to the other species. While most species exhibit a general decline in effective population size during the Mid-Pleistocene glaciations and the onset of the last glacial period, a few—such as *Asclepias syriaca, Cynanchum wilfordii*, and *Marsdenia floribunda*—demonstrate signs of recovery from these bottlenecks, suggesting higher adaptive capacity marked by early *Ne* stabilisation. In contrast, the remaining species show limited or no such recovery in effective population size.

In Asia, *C. wilfordii* and *A. venetum* have N_e_ curves with unimodal and bimodal peaks, respectively, coinciding around ∼2Ma, while *R. serpentina* shows a distinct peak at ∼200 KYA, followed by a gradual decline and a secondary peak near ∼30 KYA after which N_e_ continues to decline till ∼10 KYA. *A. pictum* depicts a single peak of N_e_ at around ∼4Ma, following a sustained decline till ∼150 KYA, followed by an immediate stabilization of N_e_. Among plants endemic to Madagascar, *C. roseus* shows an intense unimodal peak at around 900 KYA. *P. lamerei* shows notable fluctuations in N_e_ over time, with distinct peaks observed during both the Pliocene and Pleistocene glacial periods. *M. floribunda* shows bimodal peaks in N_e_ with a major peak lying in the early Pleistocene period at around ∼2Ma, followed by contraction and partial recovery at the MIS5 stage. In regions of Central America and Canada, all species peak in N_e_ at around ∼1Ma. *A. incarnata* curve depicts population expansion starting at ∼2Ma with effective population size peaking at ∼1Ma, followed by a sharp decline in N_e_ a few thousand years later. *E. panduratus*, on the other hand, shows bimodal peaks with increasing amplitudes at ∼1Ma and 250 KYA. In contrast, *A. syriaca* exhibits bimodal peaks; however, compared to the other two, the overall distribution of effective population size throughout history is undramatic and remains stabilised, especially in the earlier time intervals from around ∼500 KYA.

Overall, all the lineages show a decline in effective population size in the recent time intervals. From approximately 50 KYA, the lineages show relative stabilisation with a slight downward trend. All species exhibit high variance in N_e_ among bootstrap replicates at recent times compared to the rest of the curve. Such patterns are expected, as in recent time intervals, fewer recombination events are created with very recent coalescent histories.

## Discussion

In this paper, we report a de novo genome assembly for *Sarpgandha, R. serpentina*. As the first genome assembly of a *Rauvolfia* species native to India, this study fills a significant gap in the genomic resources available for the *Apocynaceae* family and was paleodemographically analysed in conjunction with nine other species to help understand the historical population dynamics of *Apocynaceae* and the climatic and ecological events that have shaped their evolutionary trajectories.

The demographic histories inferred from the Pairwise Sequentially Markovian Coalescent (PSMC) model revealed a general declining trend in effective population size across the *Apocynaceae* family from ∼300,000 to ∼900,000 years ago. Such a decline is most likely attributable to the intense climatic oscillations associated with the Mid-Pleistocene Transition (MPT), during which the duration of glacial cycles increased from ∼41,000 to ∼100,000 years. These climatic changes led to longer and more severe glacial–interglacial cycles, which resulted in extended periods of cold and dry conditions (Clark et al., 2006; Pisias and Moore, 1981; Schuman, 2025; van der Hammen, 1974; Verbitsky et al., 2018). Thereby, this concurrent decline in N_e_ likely reflects a shared historical bottleneck rather than an independent, species-specific demographic occurrence.

Nevertheless, species-specific traits such as dispersal ability, physiological plasticity, and biotic and abiotic interactions can modulate the response to such environmental challenges, enhancing ecological adaptation and resilience to fluctuations in temperature and precipitation. As a result, some species varied from the usual trend, depicting complex trajectories. *A. syriaca, C. wilfordii*, and *M. floribunda* displayed relatively stable Ne trajectories throughout much of the mid to late Pleistocene period, particularly during interglacial periods, from Marine Isotope Stages (MIS) 11 to 5, while *E. panduratus, P. lamerei*, and *A. venetum* demonstrated signs of recovery in *Ne* following earlier periods of decline, suggesting that interglacial intervals—characterised by warmer climates, receding ice sheets, and elevated atmospheric CO□ levels facilitated demographic expansion in some lineages. In the case of *R. serpentina*, the demographic trajectory remained relatively stable from approximately 400 KYA until a more recent decline, which may be attributed to habitat loss and anthropogenic pressures.

However, it is important to recognise that PSMC is sensitive to unaccounted substructure and sex-fluid populations; hence, peaks in N_e_ estimates may also be a result of historical subpopulation structure, as lineage subdivision can lead to a decrease in frequency of coalescent events, which can inflate N_e_ (Hilgers et al., 2025; Mather et al., 2020; Nadachowska-Brzyska et al., 2016). Therefore, sampling multiple individuals across various geographic and ecological gradients within each species can render more confidence in the demographic inferences. In addition, the amplitude of N_e_ estimates is influenced by generation time, which can vary among populations due to environmental factors such as soil properties, humidity, temperature, and precipitation. However, this limitation is largely mitigated by interpreting demographic trajectories based on their overall shape and relative trends rather than absolute amplitudes.

Overall, this study provides a comprehensive comparative perspective on the demographic history of important *Apocynaceae* species. As more genomes from this family become available, similar coalescent-based approaches can be used to infer the ecological and evolutionary processes driving interspecific and intraspecific trends in population history with greater precision. To preliminarily evaluate the utility of this genome assembly for studying monoterpene indole alkaloid (MIA) biosynthesis, several publicly available genes and transcript sequences independently isolated and involved in the biosynthetic pathways of key compounds such as ajmaline and vinorine were queried against the assembled genome. Full-length matches were successfully recovered for several enzyme-coding transcripts, including ajmaline N-methyltransferase (Accession ID: KC708445), vinorine synthase (Accession ID: AJ556780.2), vinorine hydroxylase (Accession ID: KY926696.1), and strictosidine-O-β-D-glucosidase (Accession ID: AJ302044.1).

Thus, this genome assembly can be potentially leveraged for targeted identification and characterisation of genes involved in MIA biosynthesis, helping to engineer heterologous production systems for industrial-scale synthesis of pharmacologically important indole alkaloids, thereby reducing reliance on wild plant harvesting and alleviating anthropogenic pressures. It can also prove instrumental in facilitating genetic improvement strategies to introduce resistance traits against major pathogenic threats such as *Cercospora rauvolfiae* and *Alternaria tenuis*, or in potentially modulating the yield and flowering of the plant, to further promote the conservation and sustainable utilisation of *R. serpentina*.

## Ethical Statements

This article does not contain any studies with human or animal participants.

## Acknowledgements

We thank the Ministry of Human Resource Development for the fellowship to MD. The computational analyses were performed on the Har Gobind Khorana Computational Biology cluster established and maintained by combining funds from IISER Bhopal under Grant # INST/BIO/2017/019, IYBA 2018 from the Department of Biotechnology (Grant no. BT/11/IYBA/2018/03) and ECRA (Grant no. ECR/2017/001430) from Science and Engineering Research Board (all Government of India). Funds for genome sequencing were from IISER Bhopal under Grant # INST/BIO/2017/019.

## CRediT authorship contribution statement

**Mratunjay Dwivedi**: Conceptualisation, Methodology, Writing -original draft, Formal analysis, Data curation, Investigation, Validation, Visualisation. **Nagarjun Vijay**: Funding acquisition, Conceptualisation, Supervision, Project administration, Resources, Writing - original draft, Writing - review & editing.

## Data availability

All the genome sequencing data generated as part of this project are made available through BioProject: PRJEB77109 under the experiments ERX12715045, ERX12715046, ERX12715047, and ERX12715048 on European Nucleotide Archive (ENA). The genome assembly is submitted to ENA under accession ERZ28493929.

