## Supplementary Figures for "*De novo* genome Assembly of *Rauvolfia Serpentina* and comparative paleodemographic analysis of *Apocynaceae* plants"

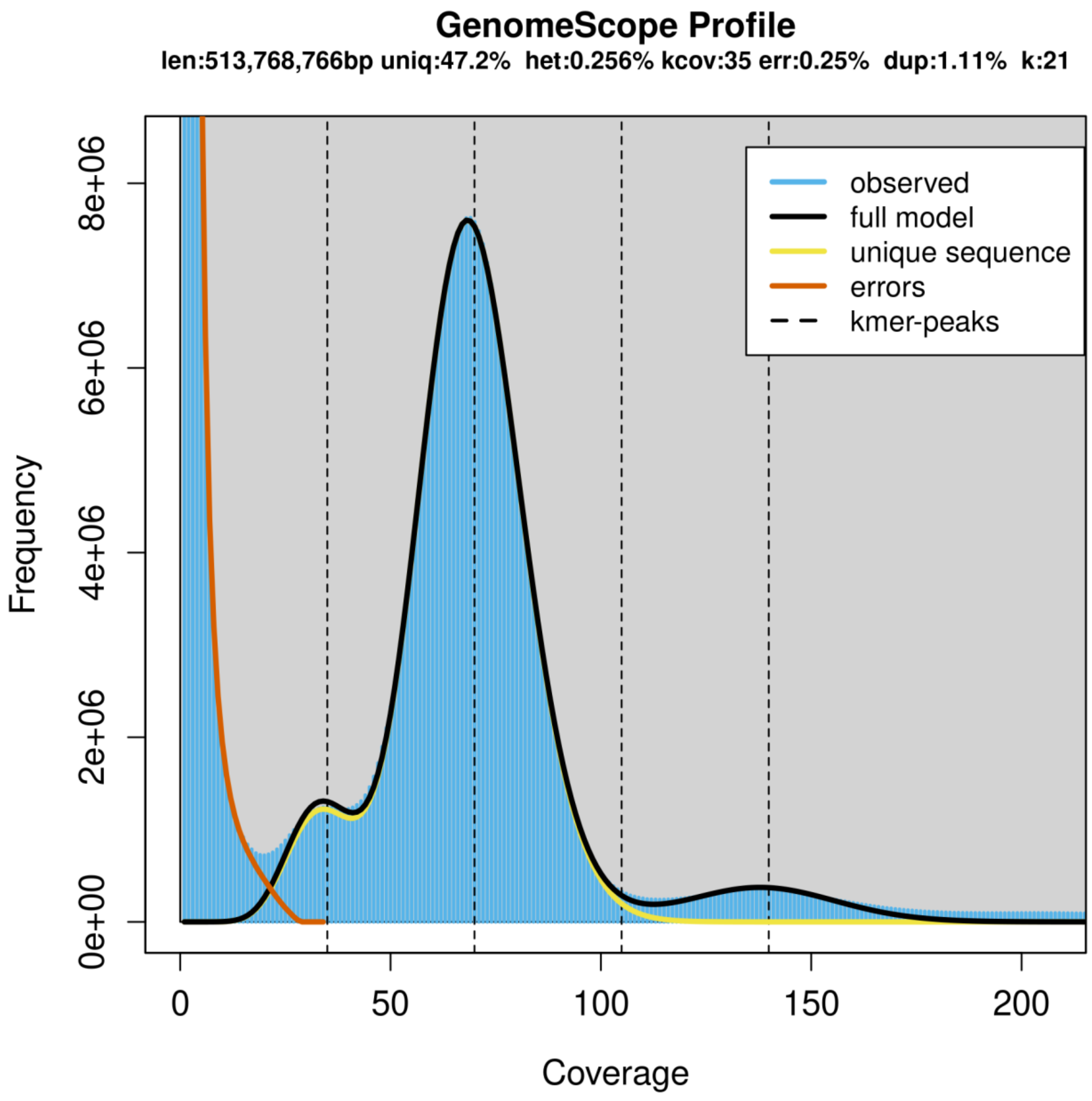


**Supplementary Fig. 1:** GenomeScope k-mer profile plot for the *R. serpentina* genome, based on 21-mers in Illumina reads. The observed k-mer frequency distribution is shown in blue, while the GenomeScope fitted model is represented by the black line. The distribution of unique k-mers is highlighted in yellow, while the putative error k-mers are plotted in red.

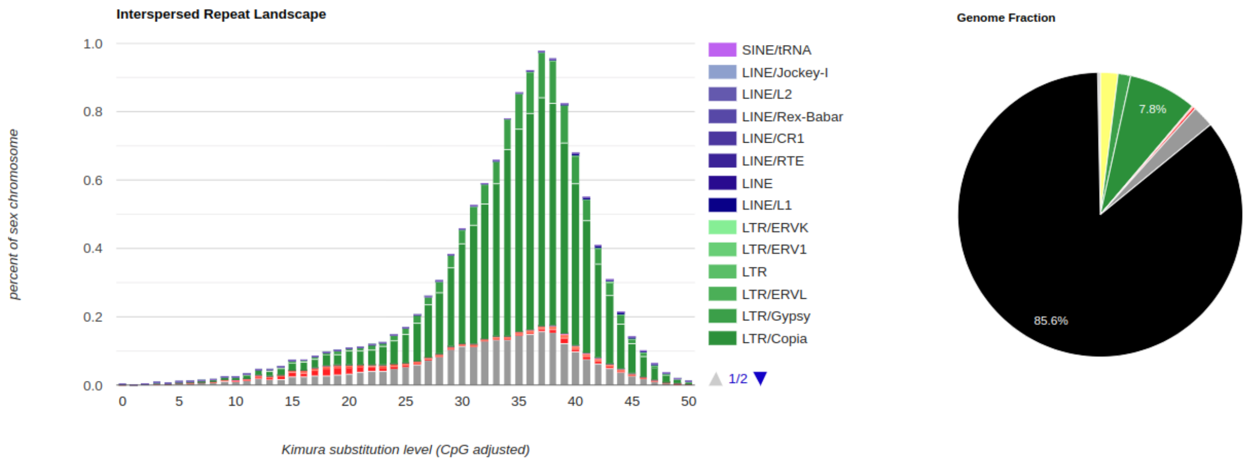


**Supplementary Fig. 2:** Analysis of repetitive elements in the *R. serpentina* genome. The repeat landscape illustrates the relative abundance of repeat classes, which are colour-coded according to the legend shown on the right. The non-repetitive fraction of the genome is represented in black.

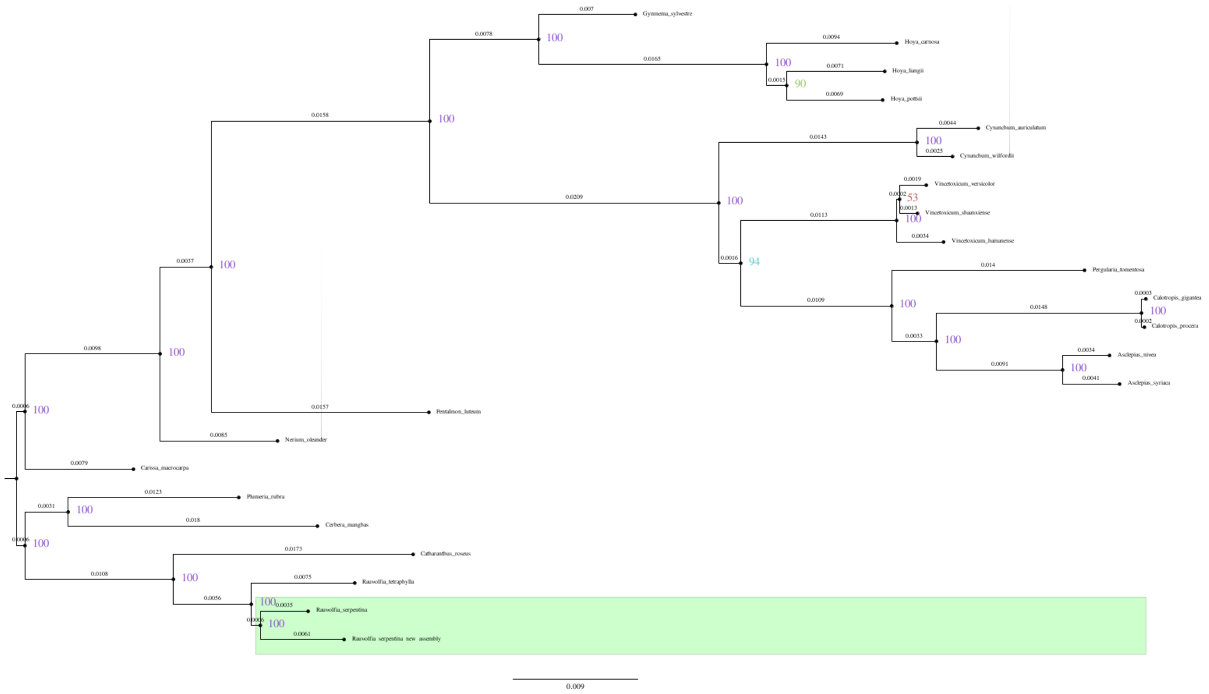


**Supplementary Fig. 3:** Phylogeny resulting from Maximum Likelihood (ML) analyses of concatenated chloroplast sequences of selected members of the family *Apocynaceae*. Numbers associated with each node represent bootstrap values inferred after 1000 maximum likelihood pseudo-replicates. The novel *Apocynaceae* sequence characterised in this study is labelled as "*Rauvolfia serpentina* new assembly" at the bottom of the tree.

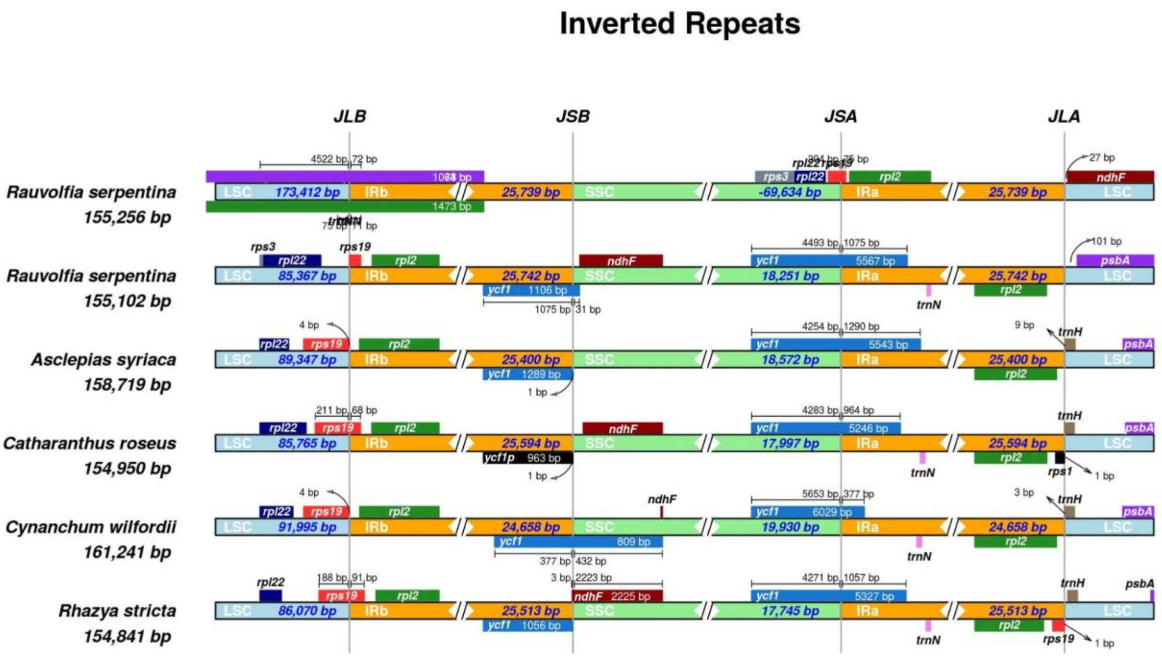


**Supplementary Fig. 4:** Comparisons of the borders of large single copy (LSC), small single copy (SSC), and inverted repeat regions (IR) among 6 *Apocynaceae* chloroplast genomes (*R. serpentina* new assembly, NC_047244.1, NC_022432.1, NC_021423.1, NC_029459.1, NC_024292.1, from top to bottom).
